# Transinfection of buffalo flies (*Haematobia exigua*) with *Wolbachia* and effect on host biology

**DOI:** 10.1101/867093

**Authors:** Mukund Madhav, Geoff Brown, Jess A.T Morgan, Sassan Asgari, Elizabeth A. McGraw, Peter James

**Author notes:** **Corresponding Author:** Peter James.

## Abstract

A widespread insect endosymbiont *Wolbachia* is currently of much interest for use in novel strategies for the control of insect pests and blocking transmission of insect-vectored diseases. *Wolbachia*-induced effects can vary from beneficial to detrimental depending on host biology and the genetic background of the infecting strains. As a first step towards investigating the potential of *Wolbachia* for use in the biocontrol of buffalo flies (BF), embryos, pupae, and adult female BF were injected with three different *Wolbachia* strains (*w*AlbB, wMel and *w*MelPop). BF eggs were not easily injected because of their tough outer chorion and embryos were frequently damaged resulting in less than 1% hatch rate of microinjected eggs. No *Wolbachia* infection was recorded in flies successfully reared from injected eggs. Adult and pupal injection gave a much higher survival rate and resulted in somatic infection and germinal tissue infection in surviving flies with transmission to the succeeding generations on a number of occasions. Investigations of infection dynamics in flies from injected pupae confirmed that *Wolbachia* were increasing in numbers in BF somatic tissues and ovarian infections were confirmed with *w*Mel and *w*MelPop in some instances, though not with *w*AlbB. Measurement of fitness traits indicated reduced longevity, decreased and delayed adult emergence, and reduced fecundity in *Wolbachia*-infected flies in comparison to mock-injected flies. Furthermore, fitness effects varied according to the *Wolbachia* strain injected with most marked reductions seen in the *w*MelPop-injected flies and least severe effects seen with the *w*AlbB strain.

## Introduction

Buffalo flies (BF), *Haematobia exigua* are obligate hematophagous ectoparasites of cattle [1]. They are present in the Australasian, Oriental and Palearctic regions of the world [2]. Both female and male BF feed 20-40 times a day on cattle, and the females only leave cattle to oviposit in freshly deposited cattle manure [3]. Their blood-feeding habits result in significant economic losses by reducing milk and meat production and causing defects in cattle leather [4, 5]. Further, BF infestation is a significant welfare issue with biting by flies causing severe irritation and, in association with a filarial nematode transmitted by BF (*Stephanofilaria* sp.), the development of lesions that range from dry, hyperkeratotic and alopecic areas to open suppurating ulcerated sores. BF are tropical and subtropical in their distribution and are mainly pests of cattle in the northern parts of Australia [6]. However, aided by a warming climate and reduced efficiency of control because of the development of chemical resistance, they have been steadily expanding their range southward [2, 6-8].

*Wolbachia*, are maternally inherited endosymbionts of insects, that are of much interest for use in the biological control of pests, most particularly as a basis for area-wide integrated control strategies for a range of insect species [9-11]. *Wolbachia* has been used in insect control programs in two main ways. First, it has been used as a means to achieve population replacement, where *Wolbachia*-infected insects impart unique characteristics such as pathogen blocking or fitness deficits, and second, by the incompatible insect technique (IIT) in which *Wolbachia*-infected males released into the population cause the production of non-viable eggs, similar to the sterile male technique [11-14]. Both of these strategies are based on cytoplasmic incompatibility (CI) and the resultant ability of *Wolbachia* to spread though uninfected or differentially infected populations [14]. Some of the novel fitness costs induced by *Wolbachia* include decreased fecundity and male competitiveness, seen in *Anopheles stephensi* infected with *w*AlbB, lifespan reduction, egg mortality, delayed larval development and altered feeding behaviour seen in *Aedes aegypti* infected with *w*MelPop [15-20].

The first successful field trial of the *Wolbachia-*based IIT technique was in Myanmar in early 1960’s to eliminate a native population of *Culex quinquefasciatus* mosquitoes responsible for transmitting filariasis [21]. Following the trial success, this strategy has been widely studied in mosquito species including *Aedes polynesiensis, Aedes albopictus, Anopheles stephensi, Culex pipiens pallens*, and in tsetse flies (*Glossina morsitans)* [10, 22-26]. Presently, *w*Mel-infected *Ae. aegypti* mosquitoes are being released in Australia, Asia (Fiji, India, Sri Lanka, and Vietnam), North America (Mexico), and South America (Colombia, Brazil) to suppress mosquito-transmitted diseases of humans such as dengue fever and Zika virus [27, 28].

The first step towards developing *Wolbachia* based control programs is the establishment of *Wolbachia* transinfected lines of the target pest. The most common method used to transinfect new hosts with *Wolbachia* has been embryonic microinjection, although injection into other stages, such as adults and pupae have also given some success [14]. Of the available transinfection procedures, embryonic microinjection is mostly preferred as *Wolbachia* are directly introduced to the pole cells of pre-blastoderm embryos using a fine needle inserted at the posterior end of the egg, desirably resulting in germline and somatic cell infection. In contrast, adult injection is usually carried out into the thoracic or abdominal regions of adults where *Wolbachia* must successfully evade or overcome a number of membrane barriers and the host immune response to become established in the germinal tissues for next-generation transmission [14]. Some instances of successful use of adult microinjection to transinfect new insect strains include the transfer of *w*Mel strain to *Drosophila melanogaster, w*AlbA and *w*AlbB to *Ae. aegypti*, and *w*Ri, *w*Mel, *w*Ha, and *w*No to the leafhopper *Laodelphax striatellus* [14, 29-31].

Buffalo flies collected from twelve locations in Australia and Indonesia were negative for *Wolbachia* infection, and this has been confirmed by more recent testing in our lab (unpublished data) [32]. However, *Wolbachia* appears to be ubiquitous in closely related horn flies (*Haematobia irritans*) (HF) suggesting that BF will also be a competent host for *Wolbachia* [32-38]. In previous studies, *Wolbachia* has been mostly sourced from the egg of the infected species for microinjection purposes [14]. Nevertheless, using cell lines of the intended host artificially infected with *Wolbachia* as the donor source has been suggested as advantageous for obtaining a high density and host context adapted *Wolbachia*. Hence, we established the HIE-18 cell line from HF to adapt *w*AlbB obtained from mosquito, *w*Mel, and *w*MelPop from *Drosophila* into the *Haematobia* spp. context prior to commencing BF microinjection.

Here, we report the results of studies towards the establishment of lines of BF sustainably infected with the *w*AlbB, *w*Mel, and *w*MelPop strains of *Wolbachia* and the dynamics and kinetics of infection in microinjected flies. The results of preliminary investigations into the related physiological costs of *Wolbachia* infection on the newly infected host BF, which are critical to considerations of the potential for use in biological control programs, are also described.

## Material and Methods

### Establishment of *Wolbachia*-infected cell cultures

A non-infected *Drosophila* cell line (JW18) was infected with the *w*AlbB (JW18-*w*AlbB), *w*Mel (JW18-*w*Mel), and *w*MelPop (JW18-*w*MelPop) strains of *Wolbachia* following the protocol of Hebert et al. (2017) to first adapt them in a closely related species [39]. JW18 cell lines infected with the three strains of *Wolbachia* were cultured in a 75 cm^2^ flask in 12 ml Schneider’s medium supplemented with 10% FBS at 28 °C (Sigma Aldrich, NSW, Australia). The *Haematobia* embryonic cell line (HIE-18) maintained in our lab without the use of antibiotics were transinfected with *w*AlbB (*w*AlbB-HIE-18), *w*Mel (*w*Mel-HIE-18) and *w*MelPop (*w*MelPop-HIE-18) as above. The infected HIE-18 lines were cultured in 75 cm^2^ flasks containing 12 ml of Schneider’s medium supplemented with 10% FBS at 28°C and subcultured every 5-6 days by splitting at a ratio of 1:2 into new flasks (Sigma Aldrich, NSW, Australia).

### *Wolbachia* isolation

*Wolbachia* were isolated from the cell lines, according to Herbert et al. (2017) [21]. Briefly, *w*AlbB, *w*Mel, and *w*MelPop infected cell lines were grown in 75 cm^2^ cell culture flasks for seven days using previously noted methods. Cells were pelleted on the eighth day by spinning at 2000 x g and washed three times with SPG buffer (218 mM sucrose, 3.8 mM KH_2_PO4, 7.2 mM MK_2_HPO_4_, 4.9 mM L-glutamate, pH 7.5), sonicated on ice for two bursts of 10 sec and cellular debris was removed by spinning at 1000 × g for 10 min at 4 °C. The supernatant was passed through 50 μm and 2.7 μm acrodisc syringe filters (Eppendorf, NSW, Australia) and centrifuged at 12000 × g to pellet *Wolbachia*. Finally, the pellet was suspended in 100 μl SPG buffer and used for microinjection.

### Embryonic microinjection

Buffalo flies were held in temporary cages for 20-30 min to collect eggs of similar age. Newly laid eggs (40 - 60 min old) were arranged on double-sided sticky tape using a paintbrush and microinjected at the posterior pole of each egg with *w*AlbB (2×10^8^ bacteria/ml) using a FemtoJet microinjector system (Eppendorf, NSW, Australia). The microinjected eggs were then placed on tissue paper on the surface of artificial manure pats to hatch. After eclosion, larvae migrated into the moist manure where they fed until pupation. Pupae were separated from the manure by flotation in water on day 7 post-injection and incubated at room temperature. Flies that emerged from the puparium by day 10 were collected and separated by sex. Females that emerged from microinjected eggs were held singly with two males for mating in small cages made of transparent acrylic pipe (6 cm diameter × 15 cm height) closed with fly mesh and a membrane feeder at the top supplying cattle blood maintained at 26 °C. A 55 cm^2^ petri-dish containing moist filter paper was placed at the base of the cages for collection of eggs deposited by the flies. Females were allowed to oviposit, and the eggs were collected until the death of the flies. Dead flies were collected and tested for the presence of *Wolbachia* using real-time PCR.

### Adult microinjection

Approximately 100-150 pupae from the BF colony at the EcoScience Precinct, Brisbane, Australia were held separately from the main colony for collection of freshly emerged female flies (2-3 hrs old) for injection. The female flies were collected within 3-4 h of eclosion from the pupae, anaesthetised using CO_2_ for 30-40 s, and then 2 μl of *Wolbachia* suspension (3×10^9^ bacteria/ml) was injected into the metathorax of each fly using a handheld micro-manipulator (Burkard Scientific, London, UK) with hypodermic needles (0.24 × 33 mm). The microinjected flies (G_0_) were blood-fed and mated with male flies at the ratio of 1:1 in small cages as described above. On day three after injection, an artificial 100 g manure pat was placed onto sand at the base of each cage. Manure pats were removed every second day, and the collected eggs were reared to adults following our standard laboratory protocols. Newly hatched G_1_ female flies were mated to potentially infected males, allowed to oviposit until death and the dead G_1_ flies then tested by real-time PCR for the presence of *Wolbachia.* Depending on the results of testing, the cycle was repeated.

### Pupal microinjection

Approximately 3000-4000 eggs from colony-reared BF were incubated and the larva grown on manure to collect freshly pupated BF for microinjection (1-2 h old). Pupae were aligned on double-sided sticky tape and injected in the third last segment at the posterior end close to germinal tissue using a FemtoJet microinjector system (Eppendorf, NSW, Australia). The microinjected pupae were then placed on moist Whatman filter paper and incubated at 27°C until flies emerged. Freshly emerged flies were separated and placed in a cage with a maximum of five females and five males each. Eggs collected from each cage every day were tested for *Wolbachia* infection. Once infection was detected, female flies were separated into a separate single cage and eggs were collected for the G_1_ line until the flies died. Later, dead females were tested for the presence of *Wolbachia* using real-time PCR.

### *Wolbachia* diagnostic assay

A modified Chelex extraction protocol from Echeverria-Fonseca et al. (2015) was used for extraction of DNA from the embryonic and adult microinjected samples [40]. Briefly, flies were homogenised using a Mini-Beadbeater (Biospec products, Oklahoma, USA) for 5 min in 2 ml screw-cap vials with 2 g of glass beads (2mm) and 200 μl of buffer containing 1 × TE buffer and Chelex^®^-100 (Bio-Rad Laboratories, CA, USA). Samples were then incubated overnight at 56 °C with 10 μl of Proteinase K (20mg/ml) and dry boiled the next day for 8 min at 99.9 °C. Finally, samples were spun at 13000 × g for 15 min, and the supernatant was stored at −20 °C until tested. For pupal-injected samples and eggs, DNA was extracted using an Isolate II Genomic DNA extraction kit (Bioline, NSW, Australia). DNA was amplified with strain-specific primers using a Rotor-Gene Q machine (Qiagen, NSW, Australia) (Table 1). Reactions were run in a total of 10 μl having 5 μl PrimeTime ® Gene Expression Master Mix (IDT, VIC, Australia), 0.5 μl each of 10 μM forward and reverse primer, 0.25 μl of 5 μM probe and 3 μl of genomic DNA. Negative and positive PCR controls were run with every batch of the samples. Optimised amplification conditions for *w*Mel and *w*MelPop were 3 min at 95 °C followed by 45 cycles of 10 s at 95 °C, 15 s at 51 °C, and 15 s at 68 °C. For *w*AlbB, the optimized amplification conditions were 3 min at 95 °C followed by 45 cycles of 20 s at 94 °C, 20 s at 50 °C, and 30 s at 60 °C. To analyse the data, dynamic tube along with the slope correct was turned on, and the cycle threshold was set at 0.01. Any sample having CT score < 35 was considered positive, negative in case of no amplification or CT score equal to zero, and suspicious where CT>35.

**Table 1:**
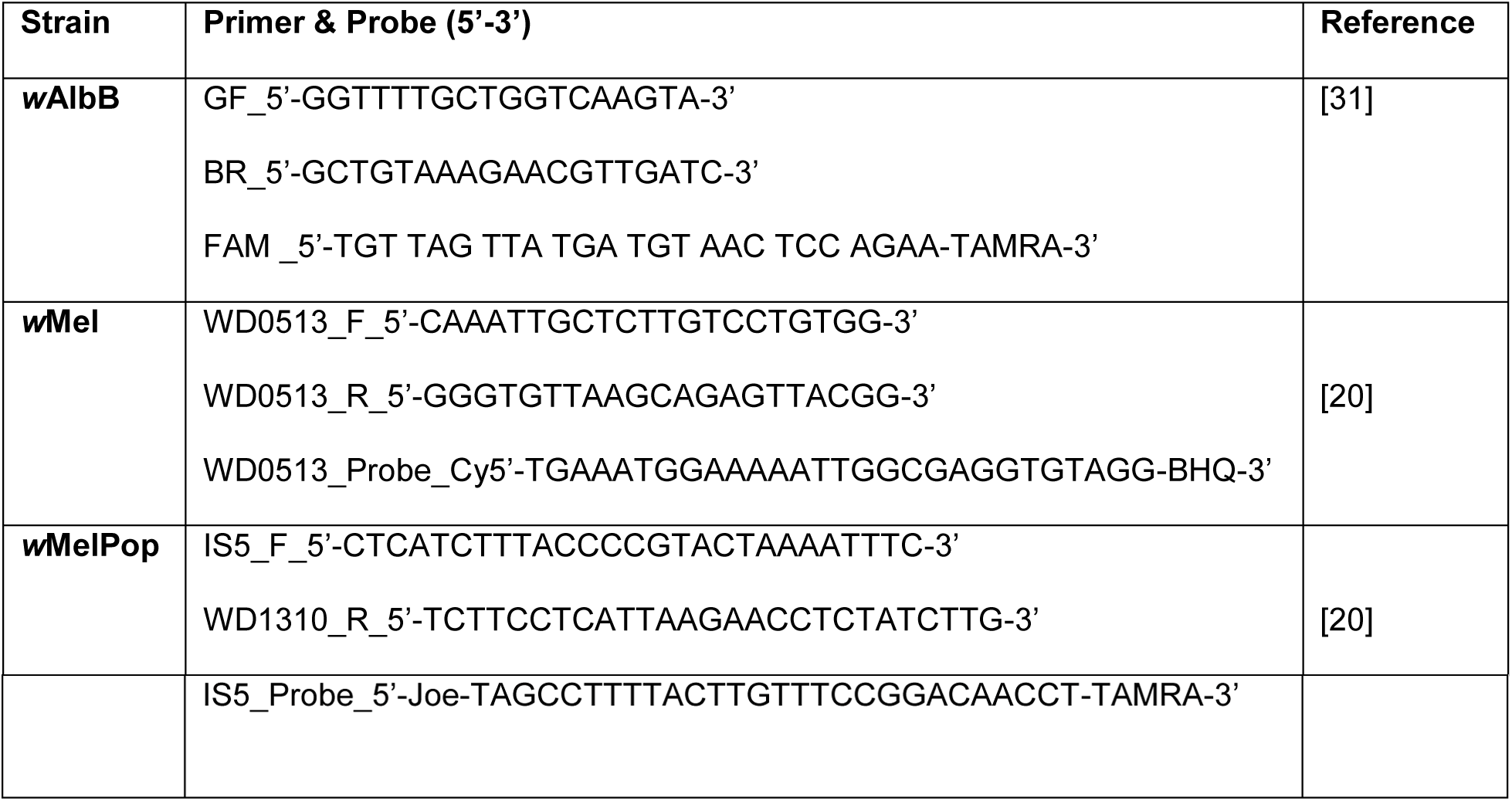
List of primers used for the *Wolbachia* Screening in the BF.

### Fluorescence *in situ* hybridisation (FISH)

FISH was carried out to visualise *Wolbachia* distribution in female BF post adult microinjection using a method slightly modified from that of Koga et al. (2009) [41]. Briefly, for the whole-mount assay, 10 BF infected with *w*Mel and *w*MelPop were collected six days post-injection and fixed in Carnoy’s solution (a mixture of chloroform, ethanol and acetic acid) at a ratio of 6:3:1 overnight. Flies were washed the next day sequentially in 100% ethanol, 80% ethanol, 70% ethanol and stored in 10% H_2_O_2_ in 100% ethanol for 30 days to quench the autofluorescence. Preserved flies were subsequently washed three times with 80% ethanol, 70% ethanol, and PBSTx (0.8% NaCl, 0.02% KCl, 0.115% Na_2_HPO4, 0.02% KH_2_PO4, 0.3% Triton X-100) and pre-hybridised with hybridisation buffer (4 X SSC, 0.2 g/ml dextran sulphate, 50% formamide, 250 μg/ml Poly A, 250 μg/ml salmon sperm DNA, 250 μg/ml tRNA, 100 mM DTT, 0.5x Denhardt’s solution) without probe two times for 15 min each. The insects were then incubated with hybridisation buffer and *Wolbachia* 16S rRNA probes overnight [42]. The next morning, samples were washed three times with PBSTx, three times for 15 min each and finally incubated in PBSTx containing DAPI (10 mg/ml) for 30 min. Samples were then rewashed with PBSTx, covered with ProLong Diamond Antifade Mountant (Thermofisher, Australia) and photographed using a confocal microscope.

### *Wolbachia* quantification assay

DNA was extracted from whole female BF post adult and pupal injection using an Isolate II Genomic DNA extraction kit (Bioline, NSW, Australia). Six flies were assayed at each point of time for determination of the relative *Wolbachia* density. Real-time PCR assays were carried out in triplicate to amplify the *Wolbachia wsp* gene [43] and host reference gene *GAPDH* (378 F_ 5’-CCGGTGGAGGCAGGAATGATGT-3’, 445 R_5’-CCACCCAAAAGACCGTTGACG-3’) on a Rotor-gene Q Instrument (Qiagen, NSW, Australia). Reactions were run in a total volume of 10 μl having 5 μl Rotor-Gene SYBR^®^ Green PCR Kit (Qiagen, NSW, Australia), 0.3 μl each of 10 μM forward and reverse primer and 2 μl of genomic DNA. Negative and positive PCR controls were included in all runs. Amplification was conducted for 5 min at 95 °C followed by 45 cycles of 10 sec at 95 °C, 15 s at 55 °C, and 15 s at 69 °C, acquiring on the green channel at the end of each step. Finally, *Wolbachia* density was calculated relative to host *GAPDH* using the delta-delta CT method [44].

### Survival assay

Two to three-hour old female adult BF were injected with *Wolbachia* (*w*AlbB, *w*Mel, and *w*MelPop) or SPG buffer (injected control) as described above and placed in triplicate cages containing ten flies each. Flies were cultured under laboratory conditions in small cages, and mortality was noted every 12 hours. Dead flies were later tested for *Wolbachia* infection individually using real-time PCR as described above. The survival assay for microinjected pupae was carried out as per the adult assay except that the number of flies in each cage was 20 (ten male and ten female).

### Adult emergence rate post pupal microinjection with *Wolbachia*

Data from five independent pupae-microinjected batches were used to analyse the effect of *Wolbachia* on adult emergence. All three *Wolbachia* strains were injected in parallel to the buffer-injected controls. The number of injected pupae varied between batches from 77 to 205 for *w*Mel, 98 to 145 for *w*AlbB, and 82 to 148 for *w*MelPop. The emergence of adults was recorded each day and the ratio of total emerged to number of injected pupae was calculated to determine the final percentage of emergence.

### Total egg production post pupal microinjection with *Wolbachia*

The effect of *Wolbachia* on the number of eggs produced by females after pupal microinjection was assessed in triplicate with ten females per cage. Buffer-injected females were used as controls and number of eggs laid and females surviving were counted every 24 hours to estimate eggs laid per day per female. Dead females were later tested for the presence of *Wolbachia* using real-time PCR.

## Results

### Embryonic microinjection of buffalo flies

Of a total of 2036 eggs microinjected with the *w*AlbB strain only 10 developed through to adult flies (six females and four males) and no infection was detected in any of the adults. Microinjecting buffalo flies is particularly difficult because of the tough chorion surrounding the egg (Fig. 1A). We observed a significant detrimental effect of injection on embryo survival and hatching (one-way ANOVA: *F*_2, 6_ = 455.3, *p*<0.0001) and identified that older eggs (40-60 min) had a better injection survival rate, 21.96% compared to 3.4% for younger eggs (10-30 min) (Tukey’s multiple comparison test: *p*=0.010) (Fig. 1B). A number of other variations of the technique were tested to improve the survival rate of eggs post microinjection. These included dechorionation of the eggs with 2.5 % sodium hypochlorite for 30 s to soften the chorion, partial desiccation to reduce hydrostatic pressure in the eggs and increase space for the retention of larger volumes of injectate, and the use of halocarbon oil (2:1 mix of halocarbon 700 and 27) to prevent desiccation of the eggs. None of these treatments markedly improved survival post microinjection (2.33%) and they also appeared to reduce egg survival in uninjected eggs (16.33%) (one-way ANOVA: *F*_2, 6_ = 181.6, *p*<0.0001) (Fig. 1C).

**Fig. 1.**
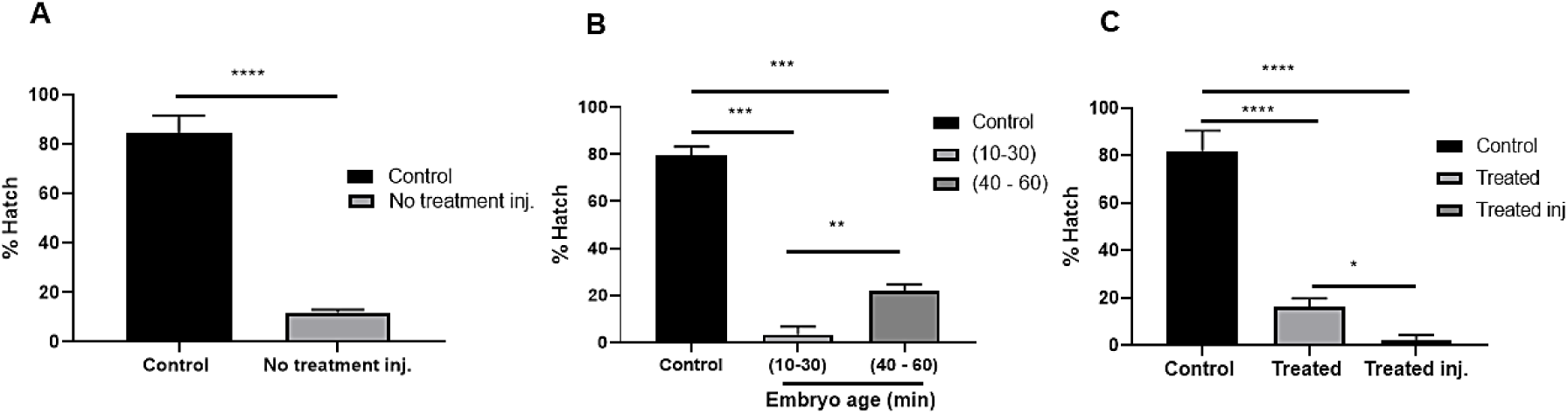
Challenges with buffalo fly embryonic microinjection. **A.** Embryonic microinjection had a detrimental effect on embryo hatching. **B.** 40-60 min old embryos survived injection better than 10 – 30 min old embryos. **C.** Eggs were dechorionated by treating with 2.5% sodium hypochlorite for 30 s and covered with 2:1 mix of halocarbon oil 700 and 27 to prevent desiccation. Eggs were sensitive to treatment and survival decreased further with the injection. Error bars are SEM. Analysis was by Student’s Unpaired t-test in (A) and Tukey’s multiple comparison test in (B) and (C); \*\*\*\**p*<0.0001.

### *Wolbachia* dynamics and tropism post adult injection

The growth kinetics of *Wolbachia* were studied in injected female flies by quantifying *Wolbachia* on days 3-11 compared to day zero (day of injection). Overall, the pattern showed an initial significant decrease in *Wolbachia* density to approximately day five followed by subsequent growth and increase in bacterial titre to day eleven in all three strains (Kruskal-Wallis test: *p*<0.0001) (Fig. 2A-C).

**Fig. 2.**
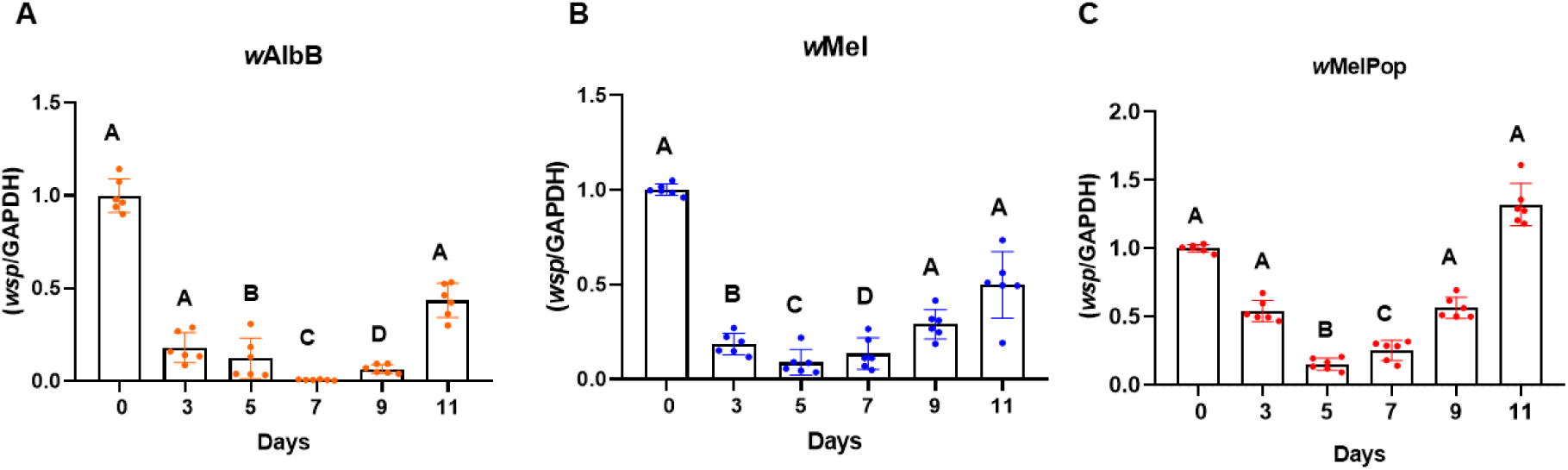
*Wolbachia* dynamics post adult microinjection of female buffalo flies assessed using real-time PCR. (A-C) *Wolbachia* dynamics measured over eleven days post-injection by analysing N = 6 for each day. Here, *Wolbachia* titre is expressed relative to the host genome. Kruskal – Wallis test and Dunn’s multiple comparison test were used to compare titres at day zero. All error bars are SEM. Bars with different letters in each graph are significantly different.

Significant variation in *Wolbachia* growth dynamics after injection required a better understanding of tissue tropism. Hence, fluorescence *in situ* hybridisation (FISH) was carried out on whole mounted BF and dissected ovaries to visualise the localisation of *w*Mel and *w*MelPop *Wolbachia* six days after injection (Fig. 3). No infection in the germline tissue was evident in any of the six samples analysed from each strain. However, *Wolbachia* was widely distributed in somatic tissues including the thoracic muscle, head, abdominal area, proboscis and legs (Fig 3).

**Fig. 3.**
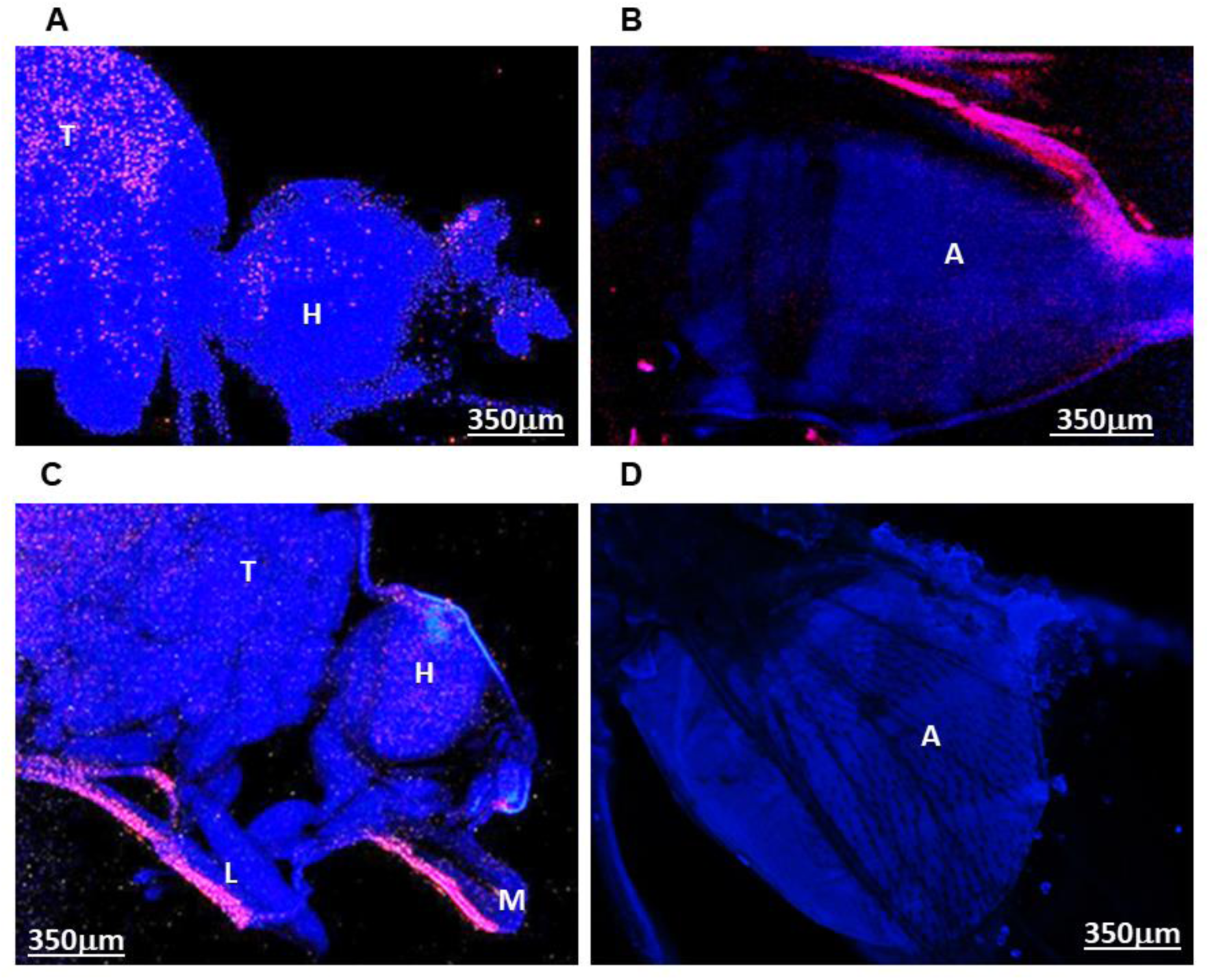
Fluorescence *in situ* hybridisation images showing localisation of *Wolbachia* six days post adult injection. *Wolbachia* is distributed throughout the BF (Blue: host, Red: *Wolbachia*). A. *w*Mel in head and thorax. B. *w*MelPop in the abdominal region. C. *w*MelPop in the head, mouthparts, thorax and leg. D. Control no probe. T: Thorax, H: Head, A: Abdomen, M: Mouthparts, L: Leg.

The PCR results for *Wolbachia* growth in flies (Fig. 2-3) suggest that the use of FISH at 6 days post-injection was too early to determine the final distribution of *Wolbachia*. Hence, we studied tissue invasion and the detailed distribution of *Wolbachia* in adult flies by real-time PCR after dissecting out the thoracic muscle, midgut, fat bodies, ovary and head at nine days post adult injection (Fig. 4A-C). *Wolbachia* were found to be replicating in all somatic tissues with *w*AlbB having an infection percentage of 33-83 % (N=6) and *w*Mel and *w*MelPop between 66-100% (N=6). No infection was found in germline tissues. However, on a few occasions first generation flies from adult injection with *w*AlbB, *w*Mel, and *w*MelPop were found positive with infection percentages of 5%, 22%, and 10% respectively, suggesting transmission via the germline tissues in these instances (see Table 2).

**Table. 2:** Summary of pupal and adult injection. G_o_ here represents injected adults and adults emerged from injected pupae. Infection was determined using real-time strain specific *Wolbachia* assays.

| Injection type | Strain | Total injected | $G_0$ (infected / total tested) (% infection) | $G_1$ (infected / total tested) (% infection) | $G_2$ (infected / total tested) (% infection) |
| --- | --- | --- | --- | --- | --- |
| Adult | wAlbB | 378<br>(19 batches) | Adult: 118/126 (95.93%) | Adult: 5/89 (5.6%) | Adult: Not tested; Egg: 0/50 (0%) |
| Adult | wMel | 441<br>(17 batches) | Adult: 117/123 (95.12%) | Adult: 27/119 (22.68%) | Adult: 0/25 (0%); Eggs: 0/100 (0%) |
| Adult | wMelPop | 417<br>(15 batches) | Adult: 103/106 (96.26%) | Adult: 10/91 (10.98 %) | Adult: 2/60 = 60 (3.3%) |
| Pupal | wAlbB | 676<br>(5 batches) | Adult : 82/90 (91.22%); Egg: 4/40 (10%) | Adult: 0/20 (0 %); Egg: 0/50 (0%) |  |
| Pupal | wMel | 820<br>(6 batches) | Adult: 82/82 (100 %) | Adult: 2/9 (22%); Egg: Not tested |  |
| Pupal | wMelPop | 741<br>(5 batches) | Adult: 88/92 (95.65 %); Egg = 0/30 (0%) | Adult: 0/23 (0%); Egg: Not tested |  |

**Fig. 4.**
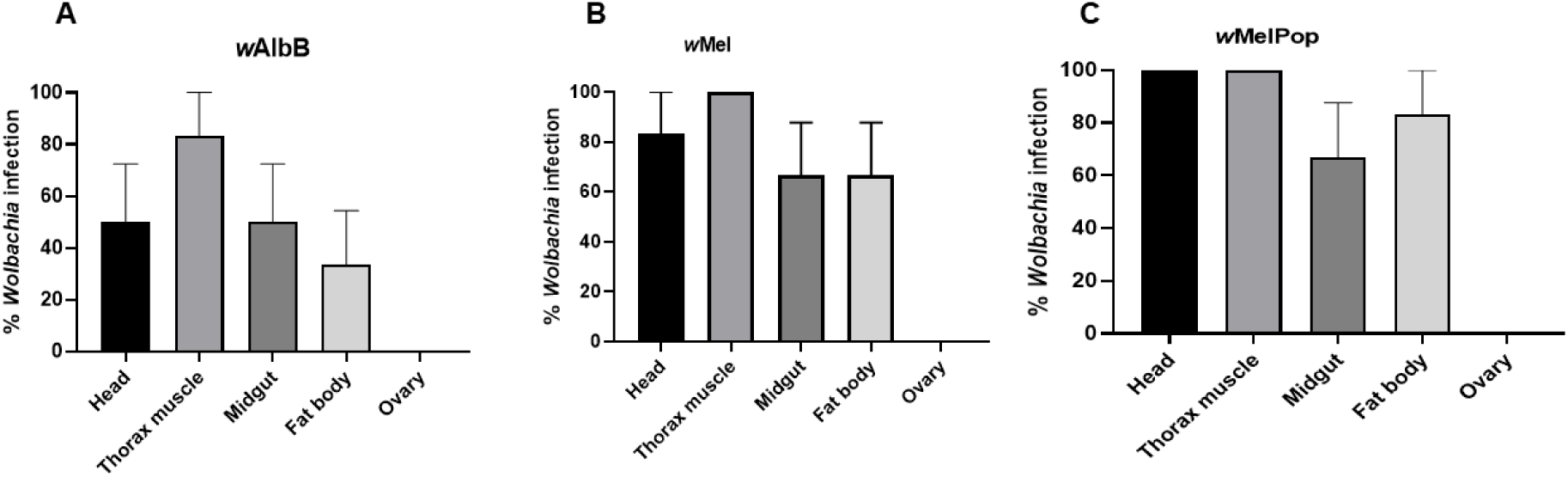
*Wolbachia* tropism post adult microinjection of female buffalo flies assessed using real-time PCR. (A-C) shows *Wolbachia* tropism in female (N = 6) nine days post adult injection. None of the *Wolbachia* strains was found in the ovaries. Bars represent SEM.

### Effect of *Wolbachia* on the survival of flies post adult injection

In order to understand the population dynamics of the flies inside the cage, survival assays were performed. The results revealed that by day seven less than 20% of the *w*MelPop and less than 50% of *w*Mel and *w*AlbB injected flies were alive (Fig. 5). Both *w*MelPop (log-rank statistic = 16.92, *p*<0.0001) and *w*Mel (log-rank statistic= 11.96, *p*=0.0005) significantly reduced longevity of female BF. However, there was no significant effect of the *w*AlbB strain in comparison to the control injected flies (log-rank statistic = 0.25, *p*=0.62).

**Fig. 5.**
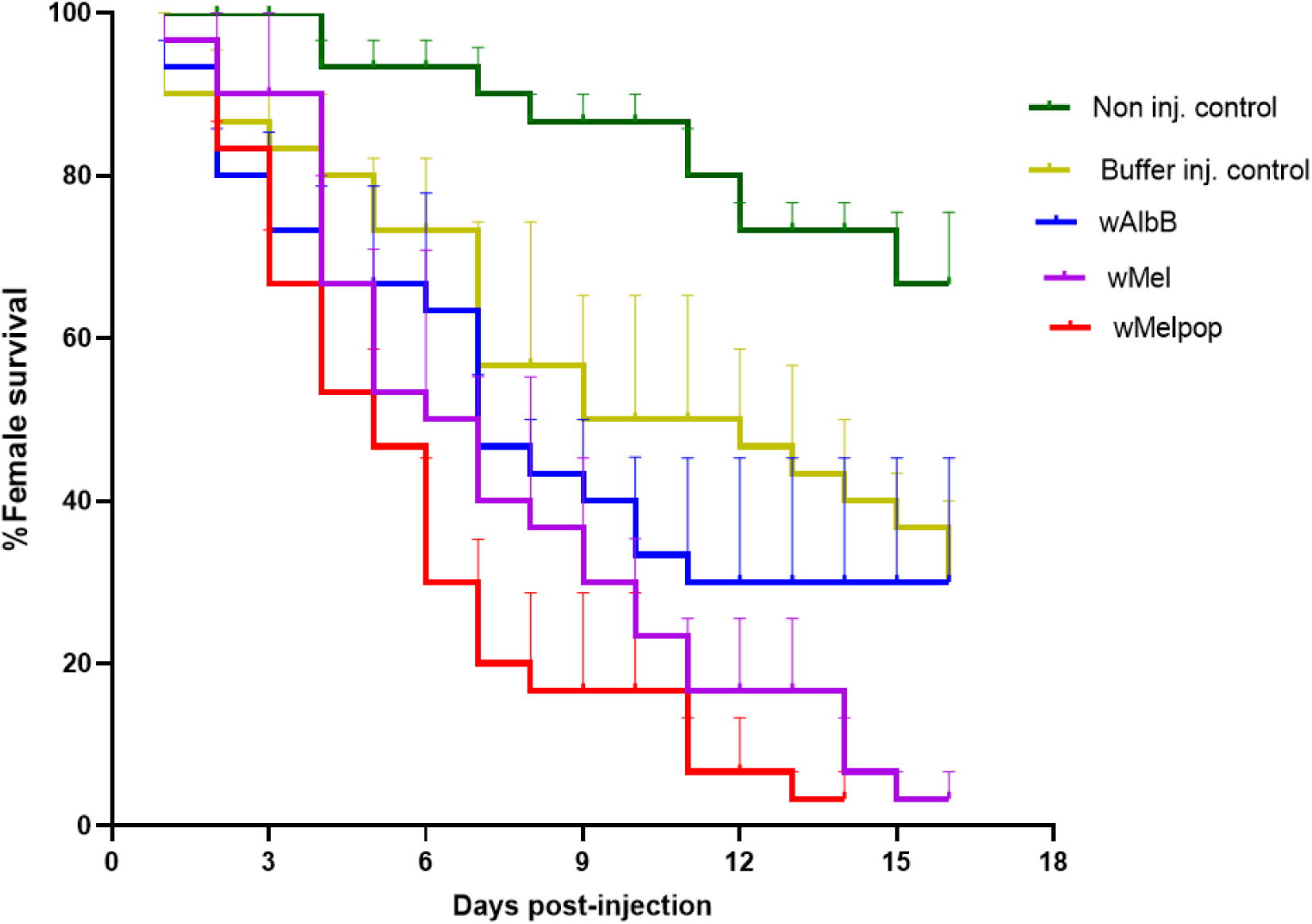
Survival of female buffalo flies post adult injection with *Wolbachia*. Triplicate cages of adult flies each containing ten females were maintained under lab culturing conditions. The number of dead flies were recorded until all died. A significant reduction in survival was observed in *w*Mel (*p*<0.0005) and *w*MelPop (*p*<0.0001) injected flies by Log-rank (Mantel-cox) tests.

### *Wolbachia* dynamics and tropism post pupal microinjection

A similar quantitative assay to that used for injected adult BF was carried out to track the dynamics and tropisms of the three *Wolbachia* strains post pupal injection. The extra time in the pupal phase resulted in 66-100% infection in the somatic tissue with *w*AlbB and *w*Mel (N=6) and 83-100% with *w*MelPop (N=6) 13 days post pupal injection (Fig. 6 A-C). Furthermore, in 16% of cases the ovaries of females injected with *w*Mel and *w*MelPop *Wolbachia* were found to be infected. Also, two first generation flies from *w*Mel-injected pupae and four eggs from *w*AlbB-injected pupae were found positive for *Wolbachia* infection (Table 2). Analysis of *Wolbachia* dynamics showed approximately the same pattern as for adult injection, where density initially decreased in the first seven days, then significantly recovered by day nine in *w*Mel (Kruskal-Wallis test: *p*<0.0001), and day 13 in *w*MelPop and *w*AlbB post pupal injection (Kruskal-Wallis test: *p*<0.0001) (Fig. 6 D-F).

**Fig. 6.**
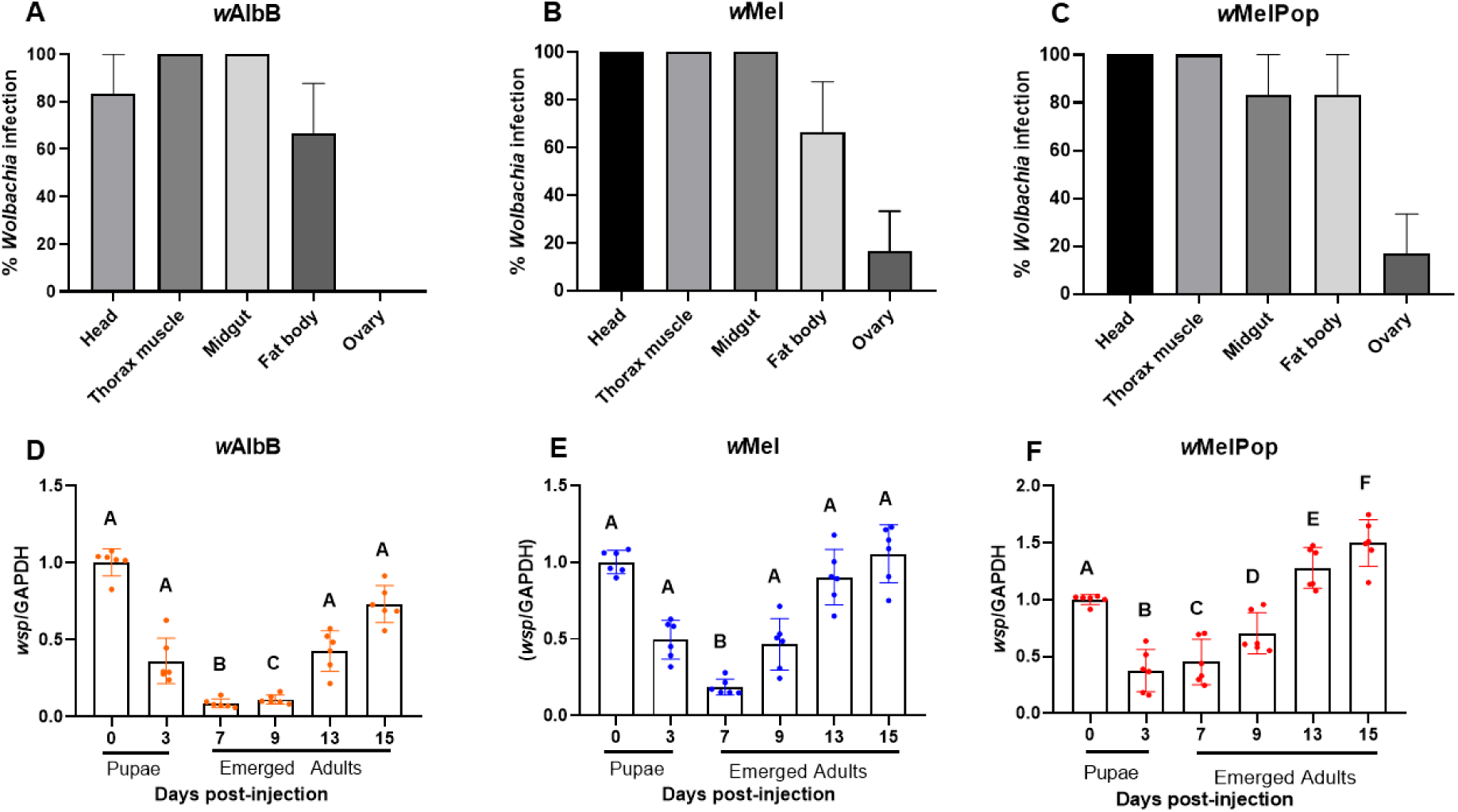
*Wolbachia* tropism and dynamics post pupal microinjection of female buffalo flies assessed using real-time PCR. A-C show *Wolbachia* tropism in female BF (N = 6) 13 days post pupal injection. Ovary infection was detected in *w*Mel, and *w*MelPop injected flies. D-F show *Wolbachia* dynamics measured over 15 days post-injection. Here, *Wolbachia* density is expressed relative to the host genome. Kruskal-Wallis and Dunn’s multiple comparison tests were used to compare titres to those at day zero. Bars with different letters are significantly different (*p*<0.05). Scale on the Y axis for wMelPop (F) is different to that for the other two strains (D,E) indicating faster growth rate with wMelPop.

### Effect of *Wolbachia* on survival of buffalo flies post pupal microinjection

A significant decrease in the longevity of BF post pupal injection was found in both sexes of *w*MelPop-injected BF (Male: log-rank statistic = 20.25, *p*<0.0001, Female: log-rank statistic =29.04, *p*<0.0001), but the effect was not significant with the two other strains (*w*AlbB: male (log-rank statistic = 2.267, *p*=0.132), female (log-rank statistic = 3.275, *p*=0.071)), *w*Mel: male (log-rank statistic = 3.027, *p*=0.1545), female (log-rank statistic = 3.467, *p*=0.063)) (Fig. 7).

**Fig. 7.**
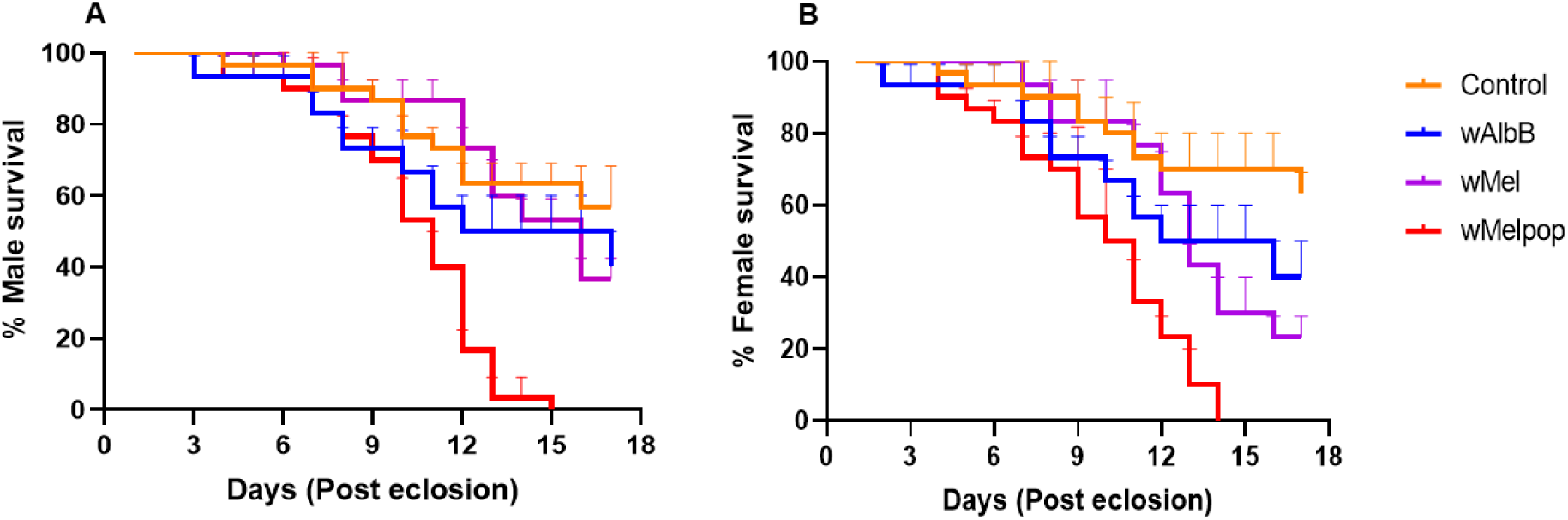
Survival of buffalo flies post pupal injection with *Wolbachia*. Triplicate cages of flies eclosed from pupae on the same day (ten males and ten females per cage) were maintained in lab culturing conditions. Mortality was recorded daily until all flies were dead. Log-rank (Mantel-cox) showed a significant reduction in the male *w*MelPop (*p*<0.0001) and female *w*MelPop (*p*< 0.0001) injected flies.

### Effect of *Wolbachia* on adult emergence rate

Infection of the somatic tissues by *Wolbachia* can have consequences on physiological processes. Non-injected control flies emerged from pupae after 3-7 days, whereas mock-injected control flies emerged from 5-7 days, *w*AlbB after 6-7 days and *w*Mel and *w*MelPop injected flies at 5-7 days post injection (Fig. 8A). It is important to note that emergence in *w*Mel and *w*MelPop injected flies was less than 2% on day 5. Overall, there was significant decrease in the percent emergence of *w*Mel (30.01 ± 3.91) (Tukey’s multiple comparison test, *p*=0.0030) and *w*MelPop (27.98 ± 3.92) (Tukey’s multiple comparison host test, *p*=0.0011) injected flies compared to the control injected flies (46.95 ± 4.15), but no significant difference was observed with the *w*AlbB-injected flies (Tukey’s multiple comparison test: *p*=0.77) (Fig. 8B). Nearly 5% of the flies that emerged from the *w*MelPop-injected pupae were too weak to completely eclose from the pupal case and had deformed wings (Fig. 8 C-D).

**Fig. 8.**
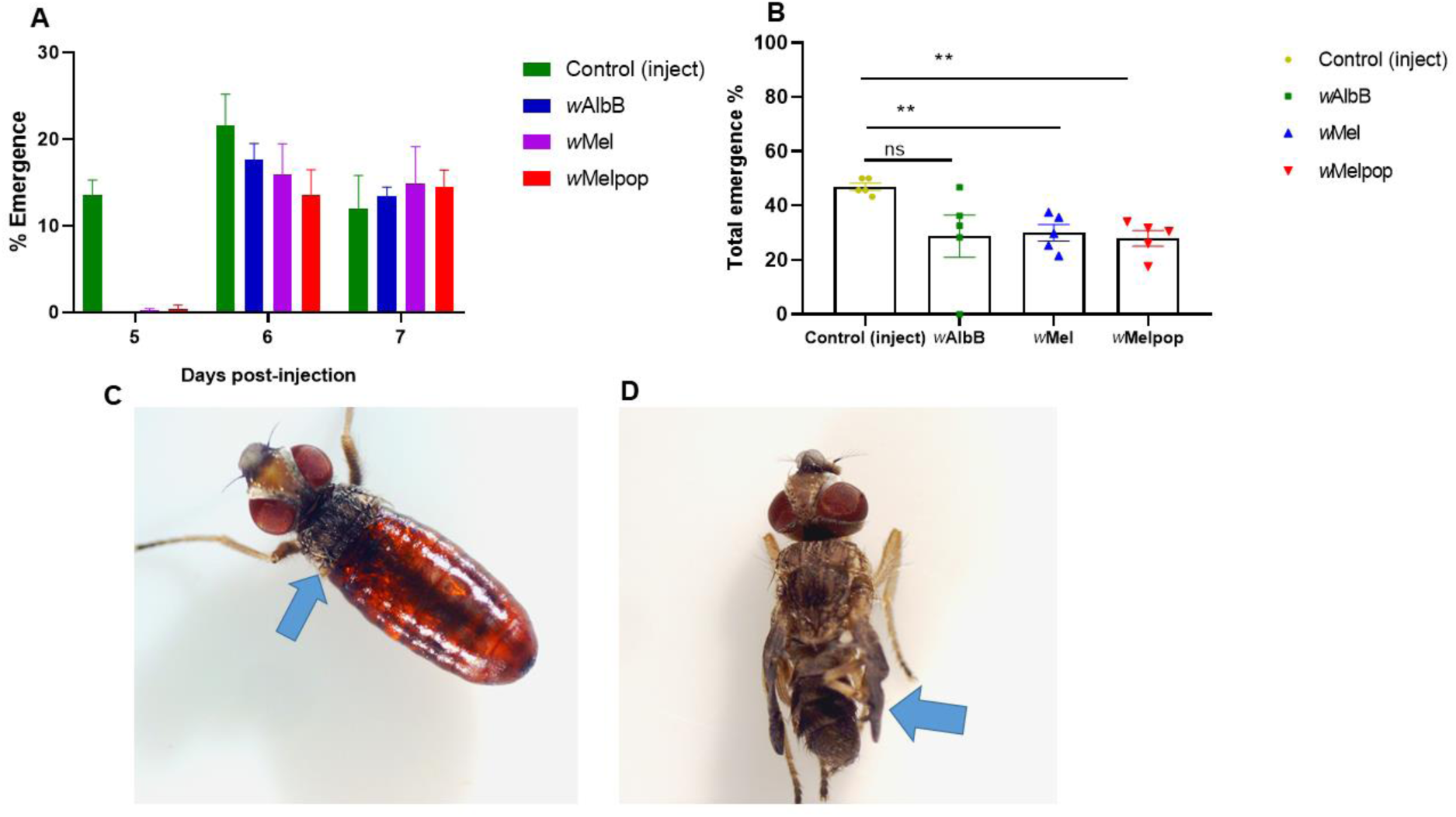
Fitness effects on buffalo fly post pupal injection with *Wolbachia*. A. *Wolbachia* delayed adult emergence. B. A significant decrease in adult emergence was observed in *w*Mel (*p*=0.0030) and *w*MelPop (*p*=0.0011) injected pupae when analysed using Tukey’s multiple comparison test. Nearly 5% of *w*MelPop flies either failed to completely eclose from the pupal case or had deformed wings.

### Effect of *Wolbachia* on egg production

Difference between infected females and non-infected females in egg production was also analysed following pupal injection with the three different strains of *Wolbachia*. Over 14 days there was a significant reduction in the total eggs laid by females infected with *w*AlbB (*p*=0.012), *w*Mel (*p*=0.0052), and *w*MelPop (*p*=0.0051) in comparison with the mock-injected flies (Fig. 9).

**Fig. 9.**
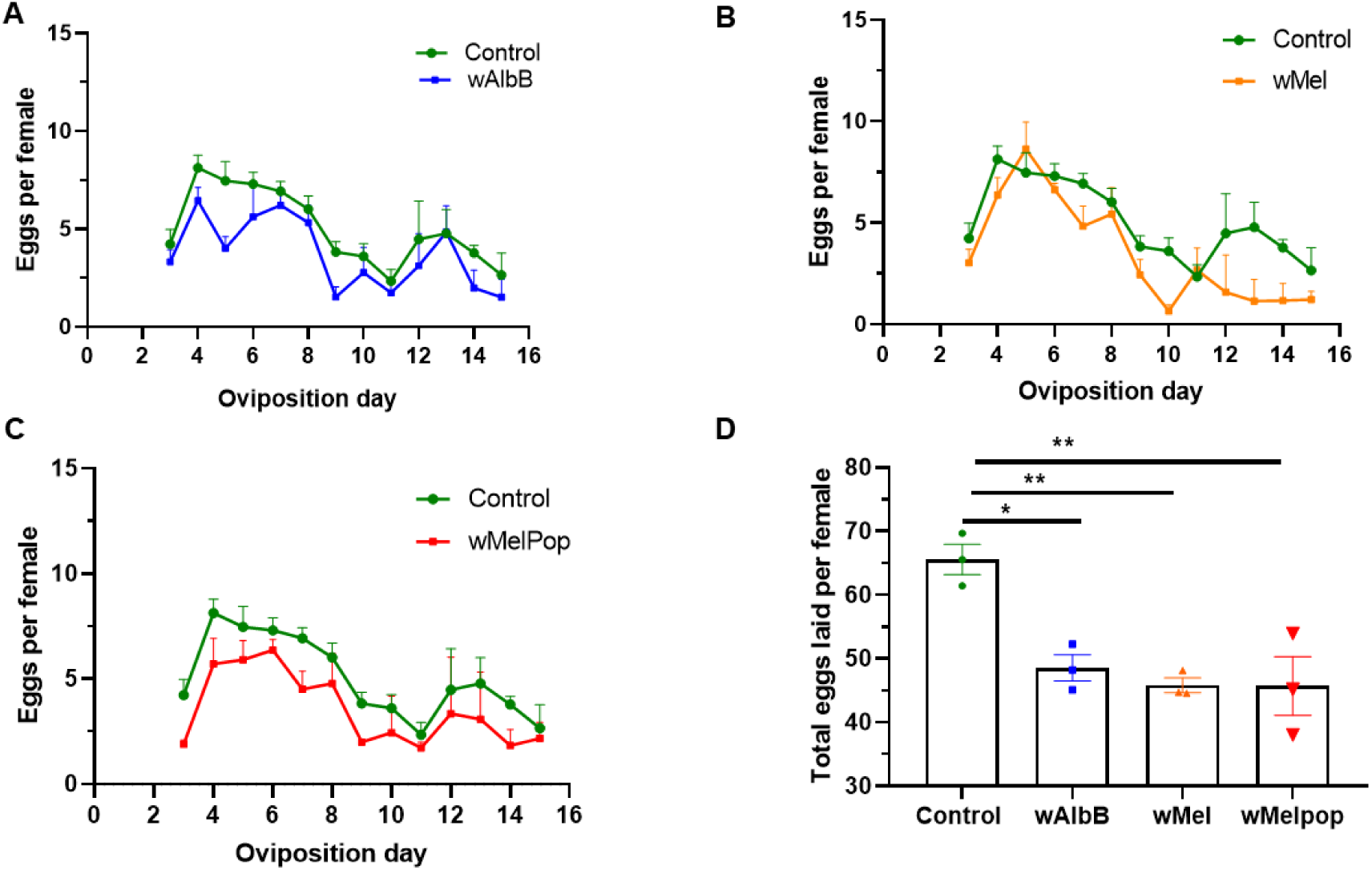
Fecundity of buffalo flies post *Wolbachia* pupal injection. Flies started laying eggs from day three post-emergence and continued until day sixteen. Eggs laid from triplicate cages each having ten females was recorded every day for (A) *w*AlbB (B) *w*Mel and (C) *w*MelPop. D. A significant difference between the total number of eggs laid per female over 13 days was found in flies infected with *w*AlbB (*p*=0.0123), *w*Mel (*p*=0.0052) and *w*MelPop (*p*=0.0051) (Tukey’s multiple comparison test).

## Discussion

Embryonic microinjection is by far the most frequently used technique to develop *Wolbachia*-transinfected insect lines, mainly because *Wolbachia* injected into the germ cells of the developing embryo provides a direct route for infection of the germ tissues in the early stage of differentiation [14]. However, this technique is also the most challenging step because the invasive procedure of egg microinjection can result in high mortality of eggs and optimal methods differ for different insect species [14, 45, 46]. Another disadvantage of this technique is that inability to determine the sex of an embryo prior to injection means that approximately half of the injected flies will be males that do not transmit *Wolbachia* to the next generation [14]. This means that many thousands of eggs must often be microinjected using specialised equipment before successful *Wolbachia* transinfection is achieved [14] and as male embryos cannot be identified, half of this effort is functionally wasted. With BF, less than 1% of more than 2000 embryos we injected subsequently hatched because the tough chorion of BF eggs caused difficulties with needle penetration, rapid blunting and high breakage rate of microinjector needles, frequent chorion tearing, and embryo damage. Treatment with sodium hypochlorite to soften the chorion, prior partial desiccation of eggs to reduce hydrostatic pressure, and the use of halocarbon oils to prevent egg desiccation during injection did not markedly improve the survival rate. Similar difficulties were experienced when attempting to use microinjection for gene transfection in closely related *Haematobia irritans* eggs. In this instance, the researchers opted to use electroporation, which is unsuitable for the introduction of bacteria [47].

Although embryonic microinjection has been the primary method used to develop transinfected insects, adult microinjection can be advantageous in that females can be selected for injection [14]. Further, adult microinjection can be performed using a simple syringe and small-bore needles delivering higher volumes of *Wolbachia* to overcome the host immunological response [14]. Our results with adult injection of *Wolbachia* were promising. Despite that injections in first few batches were made mainly with *Wolbachia* grown in *D. melanogaster* cells (*w*AlbB, *w*Mel and *w*MelPop strain), not previously adapted in *Haematobia* cells, infection rates and persistence in the injected flies were high (generally > 90%). In a few batches, transmission to the next generation was confirmed.

As oviposition by BF may begin as early as three days after eclosion from the pupae and continue until death, knowledge of *Wolbachia* distribution and dynamics in injected females was critical for us to identify the optimal timing for collecting infected eggs for the establishment of an infected colony (11-15 days). *Wolbachia* density significantly decreased to day five due to host immune response but recovered by day eleven after injection. A similar result was obtained when *w*MelPop and *w*AlbB were injected into *Anopheles gambiae* adult mosquitoes [13]. The initial host immune response was anticipated as the densities of *w*AlbB, *w*Mel, and *w*MelPop *Wolbachia* in *Haematobia* cells were also observed to initially decrease, possibly due to an innate immune response mediated by the Imd pathway (unpublished data). Real-time PCR analysis of dissected tissues nine days after injection showed *Wolbachia* to be present in all the vital somatic tissues, except for the ovarial tissues, suggesting that *Wolbachia* might need extra time to infect the ovaries. However, injection with *w*AlbB, *w*Mel and *w*MelPop *Wolbachia* caused >40% death in flies by day seven post injection, further reducing the likelihood of collecting infected eggs. Therefore, we hypothesised that microinjecting 1-2 h old pupae would give more time than with adult microinjection for *Wolbachia* to multiply, spread and establish in the ovaries. Pupal injection has previously been conducted with *Trichogramma* wasps and resulted in successful ovarian infections and persistence of *Wolbachia* in the wasp colony for 26 generations [48]. With BF, *w*Mel and *w*MelPop overcame host immune responses and established in both somatic and germline tissues. Further, in two instances, next-generation (G1) BF from *w*AlbB and *w*Mel injected pupae were positive for *Wolbachia*, indicating next-generation transmission as a result of pupal injection. The main disadvantages of pupal injection in comparison with adult injection were limitation on the volume of *Wolbachia* that could be injected and inability to distinguish female from male pupae for injection.

The *w*MelPop strain is a virulent type of *Wolbachia*, and its over replication in somatic tissues and brain cells, known in other infected insects [49, 50], may have been the reason for the early death of BF. Further, in the studies of *Wolbachia* kinetics we found a higher density of *w*MelPop than with the other two strains following both adult and pupal injection. Reduction in the longevity of infected *Ae. aegypti* mosquitoes caused by infection with *w*MelPop, decreasing the potential extrinsic incubation time for the dengue virus, was one of the characteristics that led to the hypothesis that *w*MelPop infection would reduce dengue spread [51]. Infection with *w*MelPop could also markedly reduce BF lifespan and their ability to transmit *Stephanofilaria* sp. nematodes. These nematodes have been implicated in the development of buffalo fly lesions, a significant production and welfare issue in north-Australian cattle [52]. *Stephanofilaria* has an extrinsic incubation period of up to 3 weeks in *Haematobia* spp. [53] and the life-shortening effects of *Wolbachia* shown in our study could markedly reduce the vector competency of infected flies. There is also the possibility the *Wolbachia* infection could more directly compromise the vector competency of BF for *Stephanofilaria*, as has been seen in the case another filarial nematode, *Brugia pahangi* transmitted by mosquitoes and in the case transmission of the dengue virus by *Ae. Aegypti* [54, 55].

Fecundity of insects has a significant influence on population dynamics of insect populations [56]. The successful establishment of *Wolbachia* in new host populations directly relates to the strong CI, vertical transmission and relatively more fertile egg production by infected females [57]. *Wolbachia* have been found to enhance and reduce egg production depending upon both the strain of the nematode and the host [15, 57-62]. We found that *w*AlbB, *w*Mel, and *w*MelPop significantly reduced total egg production in pupal injected flies. Also, *Wolbachia* infection caused delayed and decreased adult emergence of BF post pupal injection. *Wolbachia* being an endosymbiont lacks nutritional biosynthetic pathways and depends on its host for wide range of nutrition [63, 64]. Hence, the fitness costs observed in injected BF could be the result of competition between high density of *Wolbachia* and BF for nutritional resources such as amino acids and lipids [63, 64]. Another possibility could be that as *Wolbachia* was found in all of the critical tissues involved in the endocrine cascades for egg production and maturation in insects (midgut, neuron, fat bodies and ovary), it interfered with egg production by this means [65]. In addition, delayed larval development associated with *w*MelPop infection has been documented in mosquitoes on a number of occasions [17, 19]. If these deleterious effects are a consistent feature of *Wolbachia* infection in BF, they could have a significant impact in altering population dynamics or even crashing BF populations [17, 66]. For instance, female BF lay eggs in fresh cattle manure pats, where eggs take approximately seven days to develop into pupae depending upon the temperature and moisture content of the pat [67]. Prolonged larval development and time to eclosion of *Wolbachia*-infected BF, together with adult lifespan reduction might decrease overwintering and survival of BF, particularly during periods of unfavourable fly conditions and at the edge of the BF range.

In this work, we have shown that BF are competent hosts for the growth of *w*Mel, *w*MelPop and *w*AlbB *Wolbachia* strains and that infection can induce a number of fitness effects in the injected flies. However, embryonic injection has proven challenging with BF and to date we have not been able to establish a sustainably infected isofemale line using this technique. Pupal and adult microinjection gave much higher fly survival rates, high titres of *Wolbachia* in somatic tissues and ovarian infection and transmission to the next generation in a number of instances. Despite relatively limited testing, this gives hope for the future establishment of *Wolbachia*-infected strains of BF for the future design of *Wolbachia*-based control programs.

## ACKNOWLEDGEMENTS

We thank Prof. Scott O’Neill (Monash University, Melbourne) and the Eliminate Dengue program for the donation of the two *Wolbachia* strains *w*Mel and *w*MelPop used for this study. We also thank Dalton Baker, Dr Akila Prabhakaran, and Dr Mona Moradi Vajargah for helping with microinjection of the buffalo flies. This project was funded by Meat and Livestock Australia.

